# SC1: A Tool for Interactive Web-Based Single Cell RNA-Seq Data Analysis

**DOI:** 10.1101/2021.03.19.435534

**Authors:** Marmar Moussa, Ion I. Măndoiu

## Abstract

Single cell RNA-Seq (scRNA-Seq) is critical for studying cellular function and phenotypic heterogeneity as well as the development of tissues and tumors. Here, we present *SC1* a web-based highly interactive scRNA-Seq data analysis tool publicly accessible at https://sc1.engr.uconn.edu. The tool presents an integrated workflow for scRNA-Seq analysis, implements a novel method of selecting informative genes based on Term-Frequency Inverse-Document-Frequency (TF-IDF) scores, and provides a broad range of methods for clustering, differential expression analysis, gene enrichment, interactive visualization, and cell cycle analysis. The tool integrates other single cell omics data modalities like TCR-Seq and supports several single cell sequencing technologies. In just a few steps, researchers can generate a comprehensive analysis and gain powerful insights from their scRNA-Seq data.

## 1 Introduction

In recent years single cell technologies witnessed rapid and large scale development allowing for single cell omics data analysis to become an integral and important tool in analyzing complex biological systems and identifying heterogeneity. Clustering methods have been at the heart of scRNA-Seq analysis research Moussa & Măndoiu (2018), and currently, there are several packages for scRNA-Seq data analysis that address different aspects of the scRNA-Seq analysis challenges individually, like Monocle Trapnell et al. (2014) or Slingshot Street et al. (2018) for finding trajectories, and SCDE Kharchenko et al. (2014) for identifying differentially expressed genes. There are also software packages that implement an entire analysis workflow of scRNA-Seq data with a special focus on clustering methods or cell type identification. These packages are mostly implemented using R or Python programming languages and typically require considerable programming knowledge and effort and are not easy to use by biologists, bench scientists and researchers in life sciences in general. Notable examples of packages for scRNA-Seq analysis workflow include Seurat from Satija Lab (2015) (R), scanpy from Wolf et al. (2018) (Python), and SINCERA from Guo et al. (2015) (R).

There are also some tools available commercially accompanying single cell hardware platforms like, Loupe Browser from 10x Genomics 10x Genomics (2014), or software tools that bundle with other paid services like those provided by QIAGEN Qiagen (2020) or Immunai Immunai (2020). Although user-friendly, these software tools are not free for researchers and are to some extent a ‘black box’, with little transparency or flexibility as to the exact algorithms and parameters employed, also these tools are usually either too generalized and can not be further tailored or are too specialized to work with specific kind of data inputs. Recently, web-based scRNA-Seq analysis pipelines such as Granatum from Zhu et al. (2017) are offering a graphical interface for scRNA-Seq analysis, however, there is no widely adopted analysis workflow yet. We refer the reader to further consult recently published comprehensive reviews of single cell data methods and tools from Petegrosso et al. (2020) and Kiselev et al. (2019).

In this work, we present a web-based, highly interactive scRNA-Seq data analysis tool publicly accessible at https://sc1.engr.uconn.edu. The tool includes several data quality control (QC) options, a novel method for gene selection based on *Term-Frequency Inverse-Document-Frequency (TF-IDF)* scores Moussa & Măndoiu (2018), followed by cell type or functional clustering and visualization tools as well as differential expression (DE) analysis and gene enrichment steps. Additional analyses include various 3D interactive visualizations based on various dimensionality reduction algorithms as well as an integrated approach to clustering and ordering cells according to their cell cycle phase Moussa (2018)Moussa & Măndoiu (2020*a*). With robust default parameter values SC1 empowers researchers to generate a comprehensive initial analysis of their scRNA-Seq data in just a few steps, while also allowing them to conduct in depth interactive data exploration and parameter tuning. The process flow diagram in Fig. 1 provides a summary of the different processes implemented in SC1.

**Figure 1.**
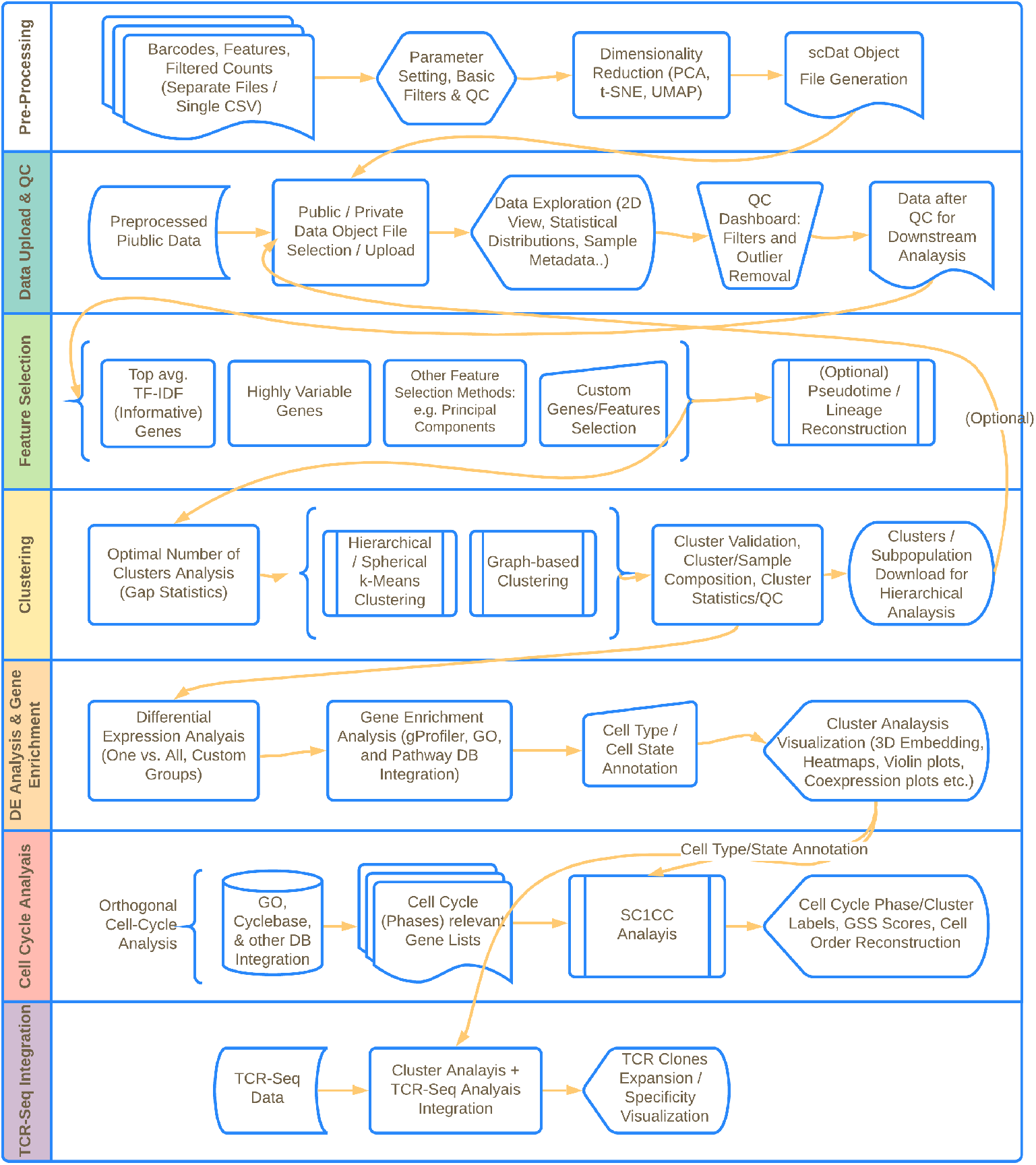
Flow diagram describing the different integrated processes of the single cell analaysis workflow implemented in SC1.

## 2 Methods

### 2.1 Datasets

To showcase the various SC1 workflow functionalities, we include examples from several publicly available datasets described in

- Fletcher et al. (2017), an olfactory stem cell scRN-Seq dataset (referred to as *OSC* dataset in the remainder of this manuscript),
- Lukowski et al. (2018), a scRNA-Seq dataset describing the detection of HPV E7 transcription at single-cell resolution in the epidermis (referred to as *HPV* dataset),
- Zheng et al. (2017), a FACS sorted combined peripheral blood mononuclear cells dataset from 10x Genomics (*PBMCs* dataset), and
- Gubin et al. (2018), a 10x Genomics scRNA-Seq set of TILs from tumor with anti-CTLA-4 treatment, describing the remodeling of the myeloid and lymphoid compartments during successful immune-checkpoint cancer therapy (*aCTLA4* dataset).

We also describe several best practices when analyzing single cell data with SC1 using the datasets described in

- Nevin et al. (2020), this 10x Genomics dataset contains the scRNA-Seq data of FACS-sorted CD3-CD19-single cells from SNS-ablated or control CT26 tumor-bearing BALB/c mice (*SNS* dataset),
- Liao et al. (2020), a 10x Genomics dataset that contains the T-cells subset extracted from the COVID-19 dataset of single cells from healthy controls and several several mildly to severely ill patients (*COVID-19* dataset), and
- Brennick et al. (2021), a recently published 10x Genomics dataset containing scRNA-Seq data of CD8+ PD-1+ TILs from mice immunized with a tumor rejection mediating neoepitope (TRMS) and a non-TRMN (*TRMNs* dataset).

The described datasets are available in SC1 data loading page as pre-processed example datasets for immediate exploration and are used in several sections of this manuscript for visualizing the different SC1 workflow steps.

### 2.2 SC1 Workflow

The SC1 workflow is implemented in the R programming language, with an interactive web-based front-end built using the Shiny framework Chang et al. (2017). In the following we present details of the main analysis steps of the workflow.

#### Data Pre-Processing

Pre-processing in SC1 tool begins with the upload of scRNA-Seq datasets in form of 10X Genomics output files (MTX and TSV files representing the filtered count matrix, features, and barcode annotations) or a single file in CSV format containing a filtered count matrix with columns representing cells and rows representing genes. Following the data upload, several pre-processing steps are carried out to prepare the data for further analysis. Starting with an initial quality control step, cells with less than a minimum number of detected genes (default 500 genes) and genes detected in less than a minimum number of cells (default 10 cells) are excluded from the dataset. Imputation is provided as a non-default pre-processing step in SC1. Imputation, aiming at inferring missing counts in single cell datasets, can be a two-edged sword that can introduce false positive signals in single cells expression signatures if not carefully applied; Empirical experiments in Moussa & Măndoiu (2019) show that over-imputation is a concern for most existing methods, hence, in SC1 we implemented the Locality Sensitive Imputation method (LSImpute) from Moussa & Măndoiu (2019), which was shown to yield high accuracy with minimum over-imputation. LSImpute implements a conservative approach to imputation in scRNA-Seq data, where only highly similar cells are used for imputation, which ensures high accuracy and minimum over-imputation. We hence provide imputation as an optional step in the pre-processing part of the analysis.

To facilitate interactive data exploration, time consuming computations such as performing dimensionality reduction of the data following basic QC are performed in the preprocessing step. SC1 pre-processing includes performing dimensionality reduction using three commonly used algorithms (see Fig. 2): Principal Component Analysis (PCA) Erichson et al. (2016), t-distributed Stochastic Neighborhood Embedding (t-SNE) projections van der Maaten & Hinton (2008), and Uniform Manifold Approximation and Projection (UMAP) McInnes et al. (2018).

**Figure 2.**
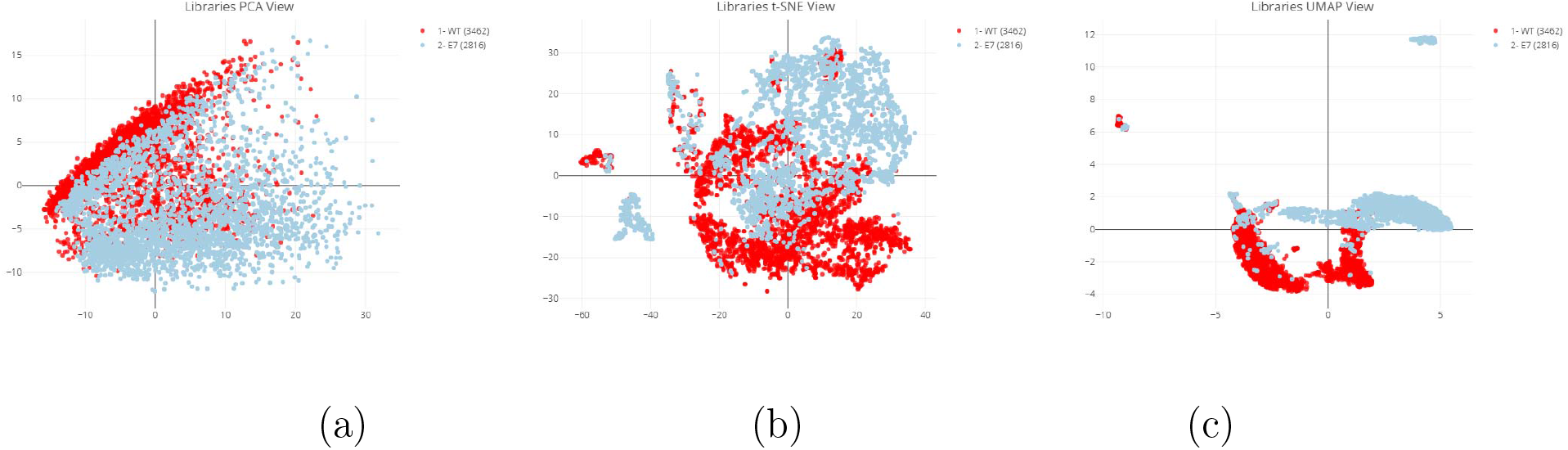
SC1: a)PCA, b)t-SNE and c)UMAP dimensionality reducing projections for the single cell RNA-Seq *HPV* dataset from Lukowski et al. (2018).

Principal Component Analysis (PCA), which historically is most commonly used, captures global variability. For larger datasets, performing matrix factorization for PCA can be a computationally expensive task and therefore an optional approach is provided: a faster but less deterministic alternative algorithm using Randomized singular value decomposition Erichson et al. (2016) can be selected in SC1 instead of the classic PCA approach.

T-distributed stochastic neighborhood embedding (t-SNE) projections are also calculated by default using the first 50 principal components. T-SNE is currently the most common nonlinear dimensionality reduction technique for single cell RNA-Seq data. The t-SNE based representation of the cells seem to avoid overcrowding while efficiently revealing locality relationship in the data.

The third approach for dimensionality reduction and data visualization is based on the recently published technique from McInnes et al. (2018): Uniform manifold approximation and projection (UMAP), which claims to preserve as much of the local and more of the global data structure than t-SNE.

#### Data Upload and Initial Exploration

At the time of this publication hundreds of thousands of single cells from experiments of different species and technologies were processed and re-analyzed with SC1 as part of the tool development; single cell technologies supported in SC1 so far include but are not limited to Smart-seq2, CEL-Seq and 10x Genomics (2014). Some of the publicly available large (file sizes exceeding 500M of 100-500K single cells) single cell RNA-seq datasets pre-processed and analyzed in SC1 are for example: the combined 10x PBMCs dataset from Zheng et al. (2017), Umbilical cord blood and bone marrow single cells from the Human Cell Atlas data portal https://preview.data.humancellatlas.org/.

Pre-processed data is saved in SC1’s “.scDat” file, an R object file that can then be uploaded for interactive analysis. Several publicly available datasets from Liao et al. (2020), Fletcher et al. (2017), Nevin et al. (2020), Lukowski et al. (2018), Zheng et al. (2017) and Gubin et al. (2018) spanning different scRNA-Seq technologies are readily provided in SC1 as pre-processed example datasets for immediate exploration and are used in this manuscript for visualizing the different SC1 workflow steps. Initial data exploration starts with ‘At-aglance’ view of the uploaded data (Fig. 3), providing basic summary statistics including the number of expressed genes and the number of cells per library, description of the loaded dataset and labels of the samples/libraries, with the ability to relabel the libraries. The two dimensional views of the data can be displayed based on the three pre-computed projections PCA, t-SNE, and UMAP.

**Figure 3.**
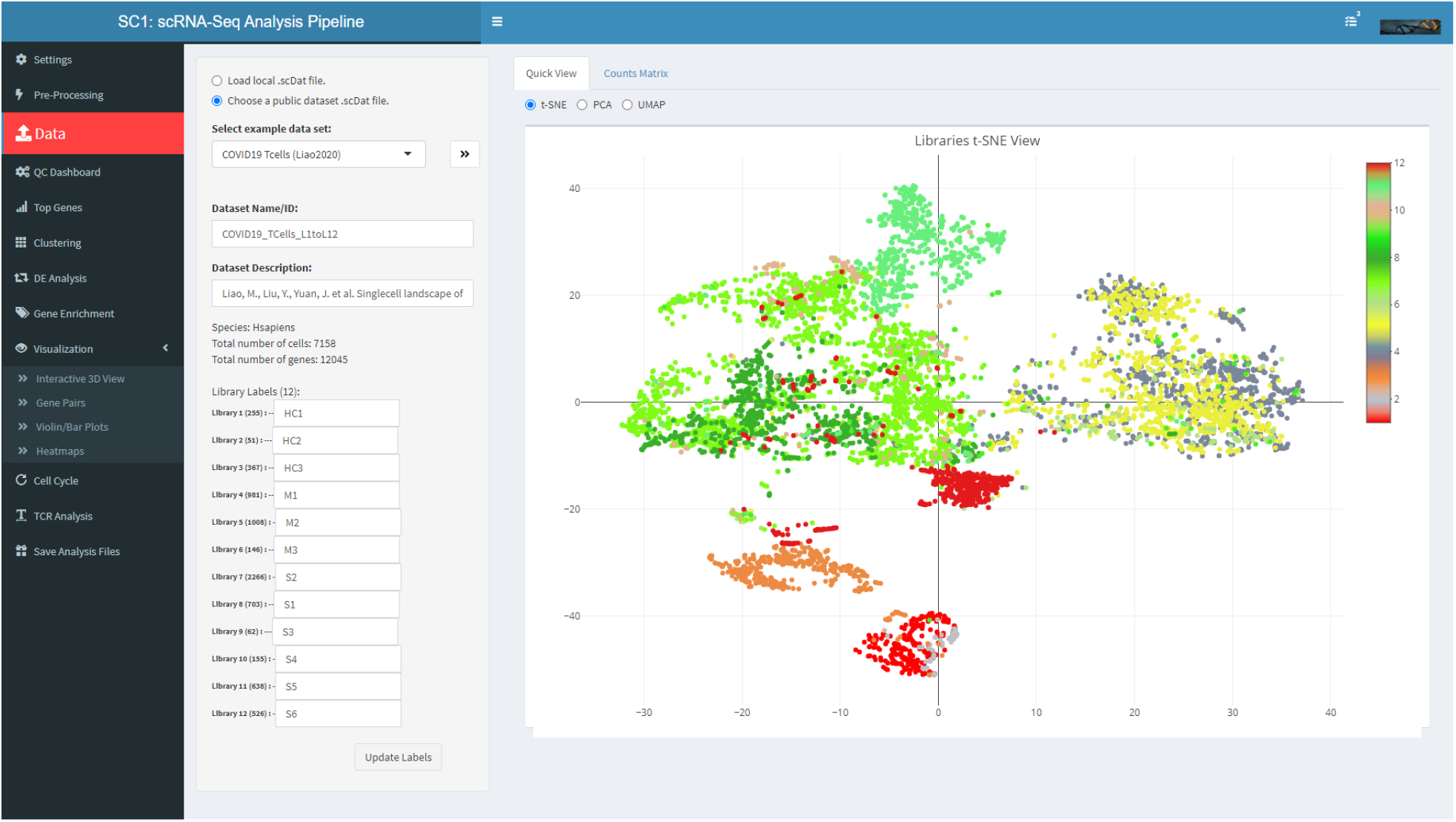
SC1 Data Upload View of the T-cells population from Liao et al. (2020) COVID-19 dataset showing 12 libraries/samples from different patients with different COVID-19 severity.

#### Quality Control Dashboard

Before further analyses, SC1 allows users to perform additional Quality Control (QC) checks as shown in Fig. 4, whereby poor quality cells and outlier cells and genes can be excluded from subsequent analysis. The tool implements widely used criteria for cell filtering: library size, number of detected genes, as well as the fraction of reads mapping to mitochondrial genes, ribosomal protein genes, or synthetic spike-ins. SC1 also allows outlier removal based on the ratio between the number of detected genes to total read/UMI count per cell.

**Figure 4.**
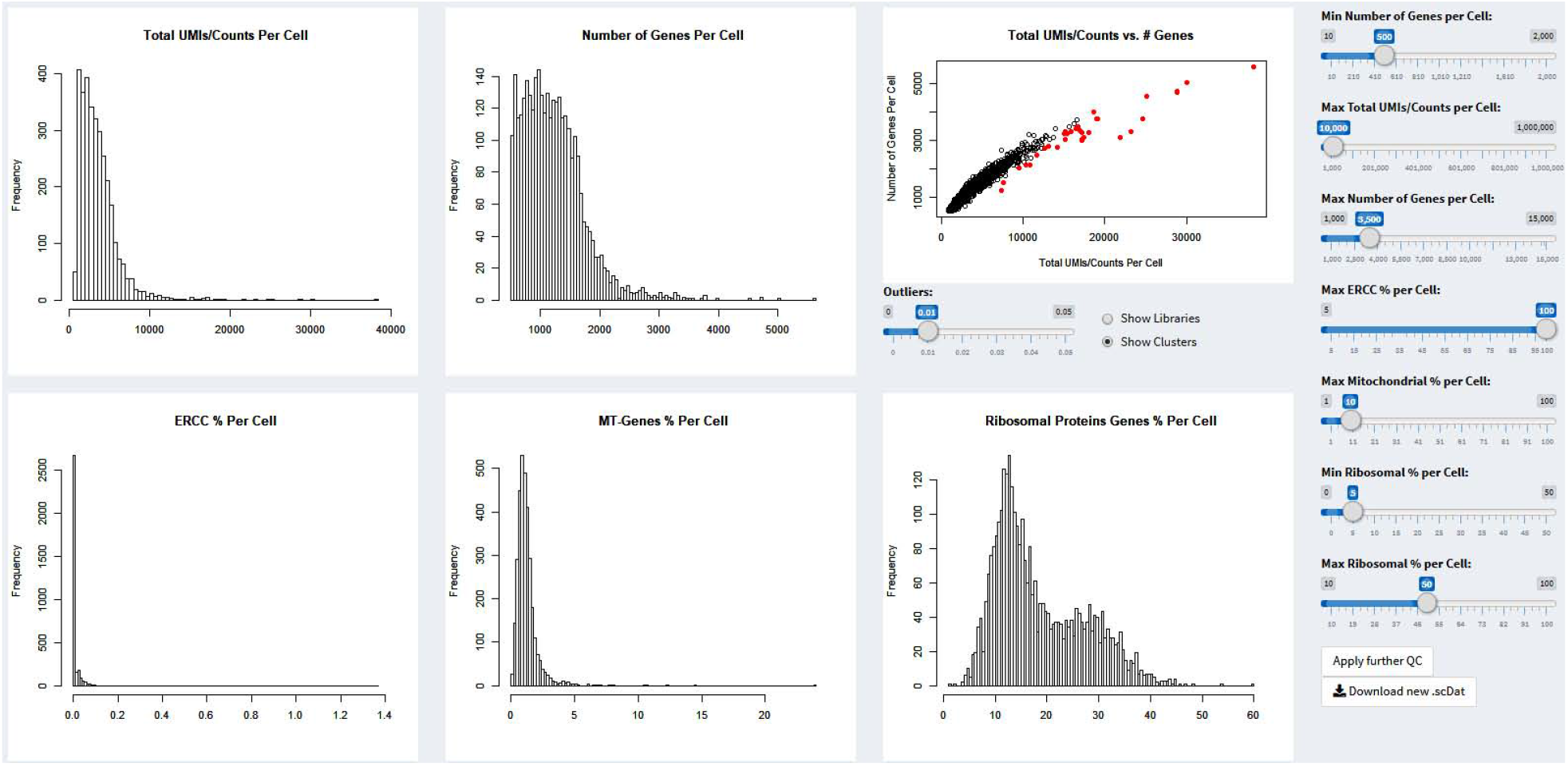
SC1 QC dashboard showing data from the *aCTLA4* dataset Gubin et al. (2018).

#### Gene Selection

SC1 implements a novel method of selecting informative genes based on the average TF-IDF (*Term Frequency times Inverse Document Frequency*) scores, as detailed in Moussa & Măndoiu (2018). To summarize, as defined in Leskovec et al. (2014), TF-IDF, a data transformation and a scoring scheme typically used in text analyses is defined as follows: Given a collection of *N* documents, and let *f*_*ij*_ be the number of occurrences of word *i* in document *j*. The *term frequency* of word *i* in document *j*, denoted by *TF*_*ij*_, is defined as

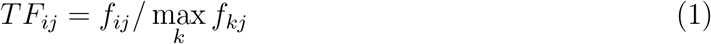

Here, the term frequency of word *i* in document *j* is the number of occurrences normalized by dividing it by the maximum number of occurrences of any word in the same document. After normalization, the most frequent word in a document always gets a term frequency value of 1, while other words get fractional values as their respective term frequencies. The *Inverse Document Frequency* of word *i* is defined as

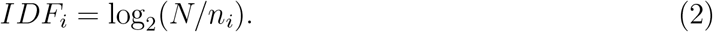

where *n*_*i*_ denotes the number documents that contain word *i* among the *N* documents in the collection. Finally, the *TF-IDF score* for word *i* in document *j* is defined to be *TF*_*ij*_ *× IDF*_*i*_. Words with the highest TF-IDF score in a document are often the terms that best characterize the topic of that document. TF-IDF scores are applied to scRNA-Seq data by considering the cells to be analogous to documents; in this analogy, genes correspond to words and UMI counts replace word counts. The TF-IDF scores can then be computed from the UMI counts (or expression values) using equations (1) and (2). Similar to document analysis, the genes with highest TF-IDF scores in a cell are expected to provide most information about the cell type. To select the genes that best separate cell populations before performing clustering and DE analysis, we estimate and fit a mixture of Gaussian models to the distribution of TF-IDF gene averages and select the genes assigned to the mixture components with highest means given a minimum number of genes as a method parameter (Fig. 5). Genes with highest average TF-IDF scores differentiate best between heterogeneous cell populations; visually this leads to a clear “chess-board” effect in the heat map constructed using the top average TF-IDF genes as shown in Fig. 6.

**Figure 5.**
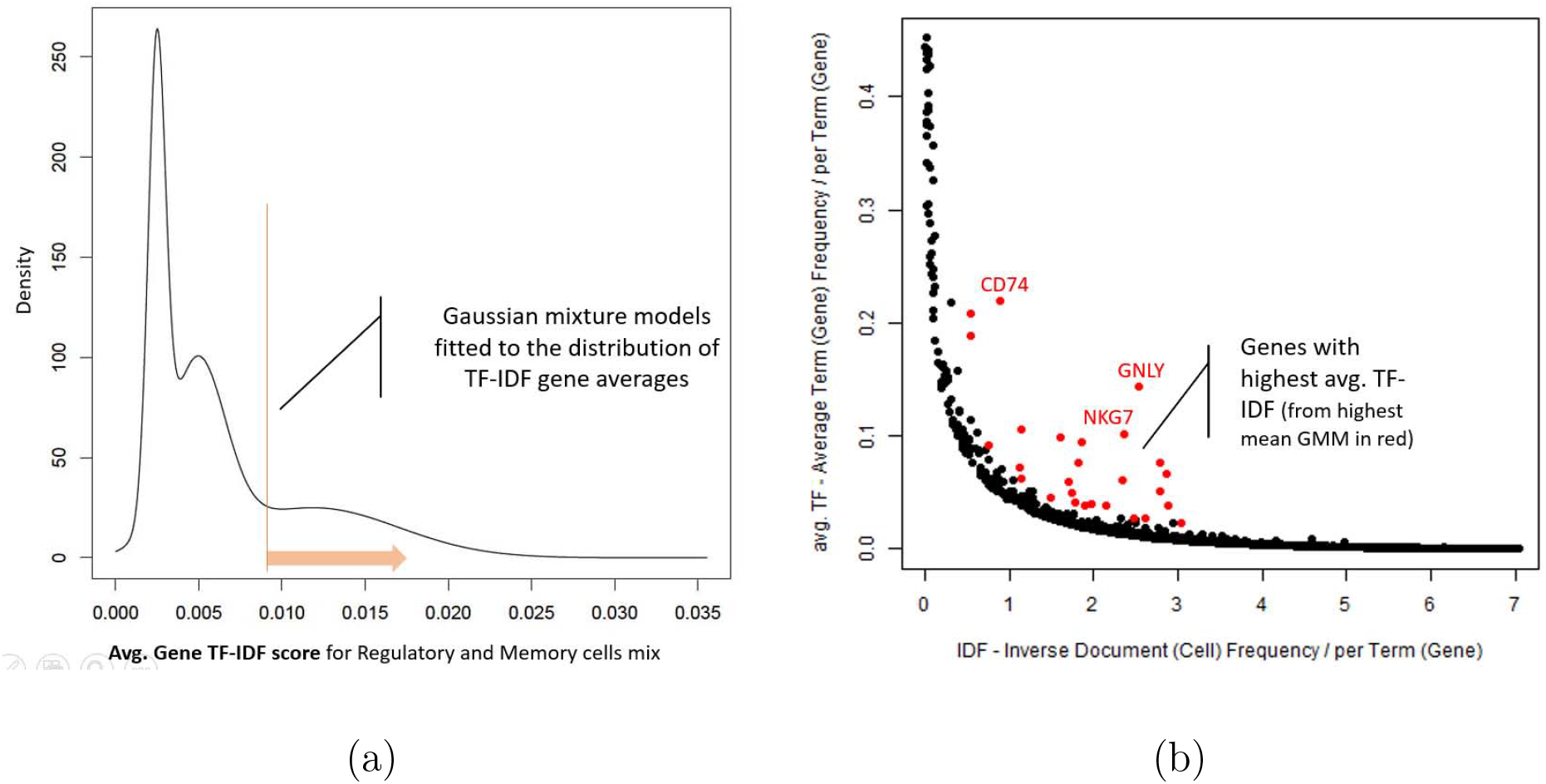
a) Gaussian mixture models fitted to genes’ average TF-IDF scores. b) Dot plot of IDF-score vs. average TF score for each gene. Genes with highest average TF-IDF scores are in red.

**Figure 6.**
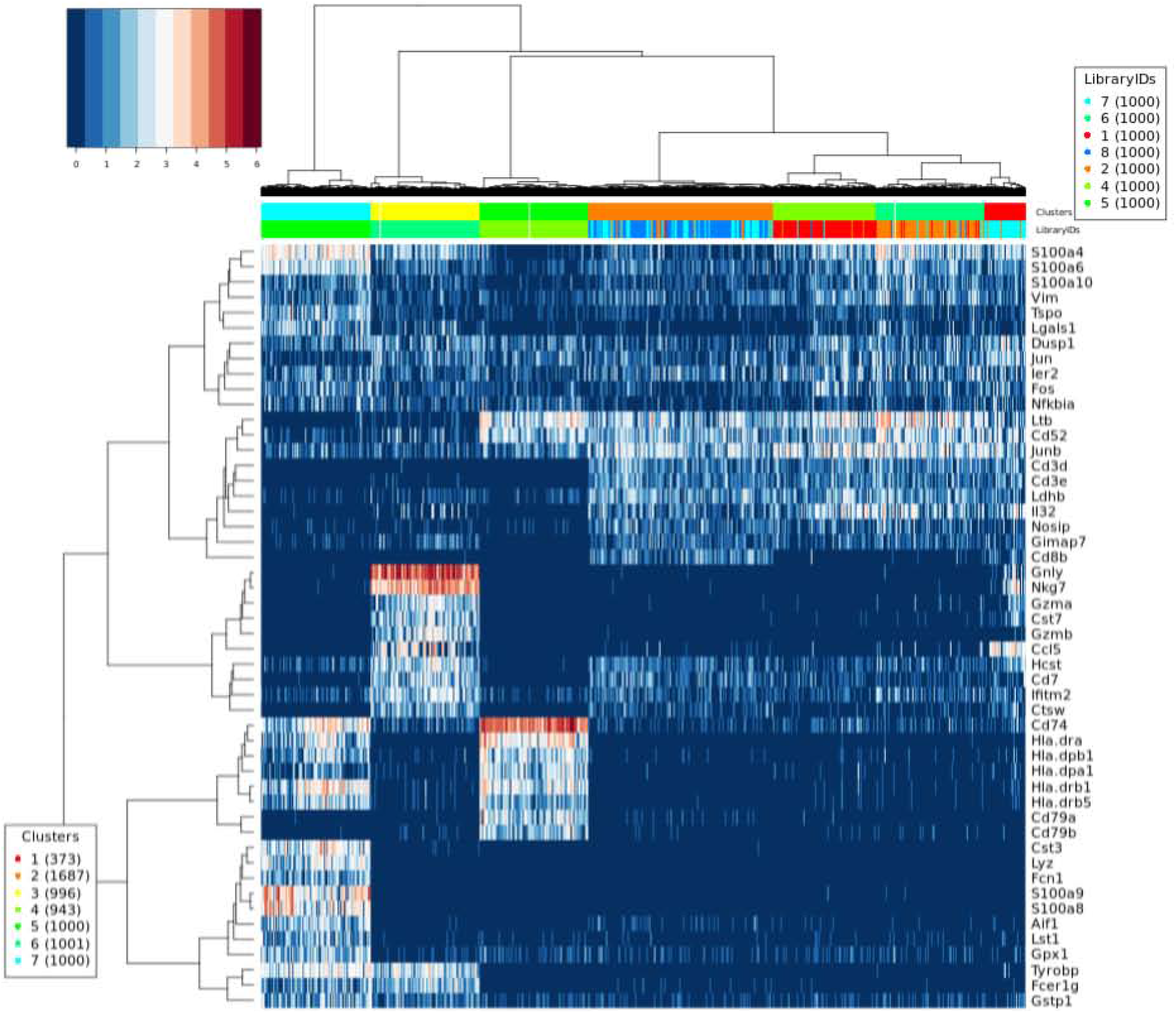
Heat map of genes with top average TF-IDF scores for cells of the 7-class *PBMCs* mixture from Moussa & Măndoiu (2018).

#### Clustering

By default, SC1 automatically infers the number of clusters using the Gap Statistics method as described in Moussa & Măndoiu (2018). However, users can also manually specify the number of clusters based on prior knowledge of the expected sample heterogeneity. Valuable insight into sample heterogeneity is also provided by inspecting the heat map generated using the top TF-IDF genes (Fig. 6) before clustering. Clustering can be performed using Ward’s Hierarchical Agglomerative Clustering or Spherical K-means (both using the top average TF-IDF genes as features) or using Graph-based Clustering using binarized TF-IDF data as described in Moussa & Măndoiu (2018). Several visualizations describe clustering details in Fig. 7.

**Figure 7.**
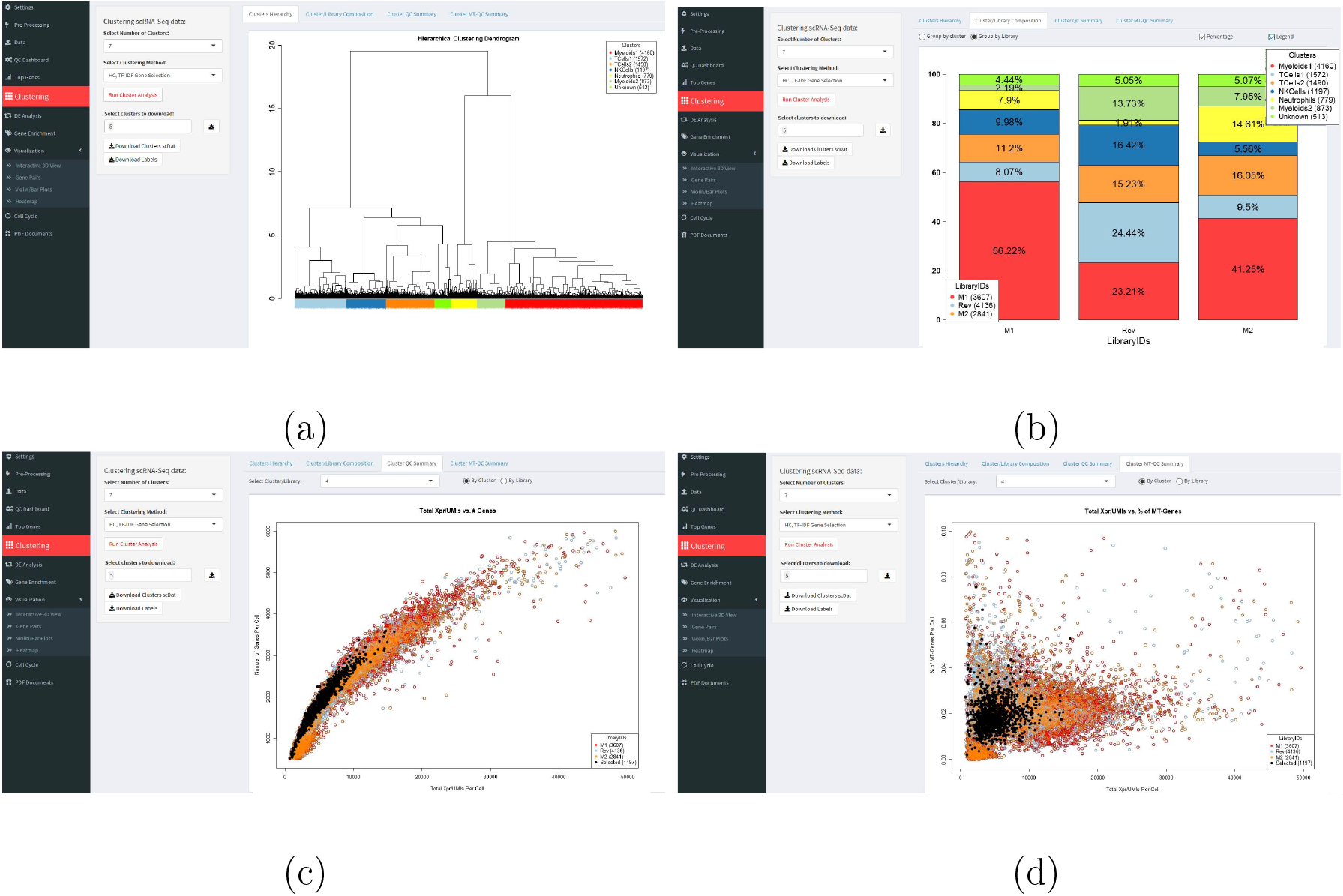
SC1 clustering analysis visualization a) Clustering results represented as a dendrogram. b) Composition bar plots specify the % of each cluster in each library and vice versa c) Scatter plot for the relation between number of detected genes and total cell size also highlights a selected cluster for QC assessment. c) Scatter plot for the total Mitochondrial content of cells in relation to total read/UMI count.

##### a) Ward’s Hierarchical Agglomerative Clustering algorithm using the top average TF-IDF genes as features

Here, hierarchical agglomerative clustering algorithm using Cosine distance is applied to the *log*2(*x* + 1) transformation of the data. Only most informative genes with top average TF-IDF scores as described in section 2.2 are used to construct the expression profiles of the cells. A dendrogram (hierarchical clustering tree structure) representing the resulting hierarchy of the cells is displayed with the tree cut at the specified number of clusters.

##### b) Spherical K-means algorithm using the top average TF-IDF genes as features

Similar to hierarchical clustering, genes with top average TF-IDF scores as described in section 2.2 are used to construct the expression profiles of the cells. The *log*2(*x* + 1) transformation of these profiles is used as input for spherical K-means clustering algorithm using Cosine distance.

##### c) Graph-based Clustering using binarized TF-IDF data transformation

This method applies Louvain graph-based clustering algorithm with the binarized version of the TF-IDF transformation of the gene-cell count matrix of the scRNA-Seq data as described in Moussa & Măndoiu (2018).

#### Lineage and Pseudo-time Inference

SC1 integrates with slingshot R package for lineage inference and pseudotime reconstruction. Using the three dimensional t-SNE presentation of the cells, the slingshot algorithm suggests the possible differentiation lineages and corresponding curves taking into account the clusters identified in a dataset as described in Street et al. (2018). In Fig. 8 we demonstrate the lineage of the OSC dataset as identified by the SC1 Slingshot package integration.

**Figure 8.**
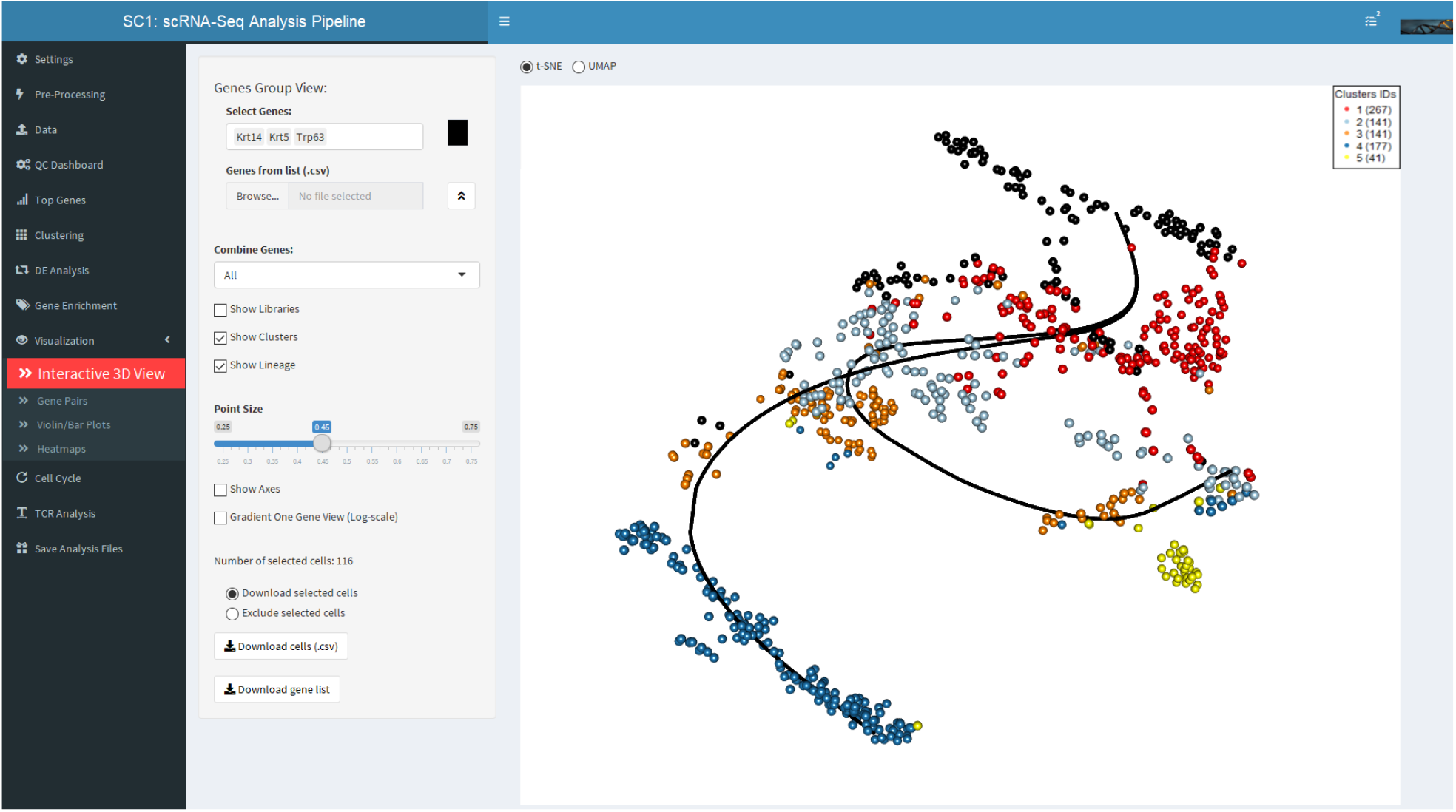
SC1 3DtSNE view of the OSC dataset showing clusters and lineage curves as identified by SC1 Slingshot package integration

#### Differential Expression Analysis

Differential expression (DE) analysis is done by performing “One vs. the Rest” t-tests for each of the identified clusters. The t-test uses the Welch (or Satterthwaite) approximation with 0.95 confidence interval by calling the t-test available in R stats package. Results of the Log2 Fold Change and the p-value from the analysis are provided as a downloadable numeric matrix (Fig. 9). A custom test of two selected groups of clusters or libraries is also provided, with results provided both as a downloadable numeric table and as a Volcano plot visualizing the Log2 Fold Change and p-values for the tested groups (Fig. 10).

**Figure 9.**
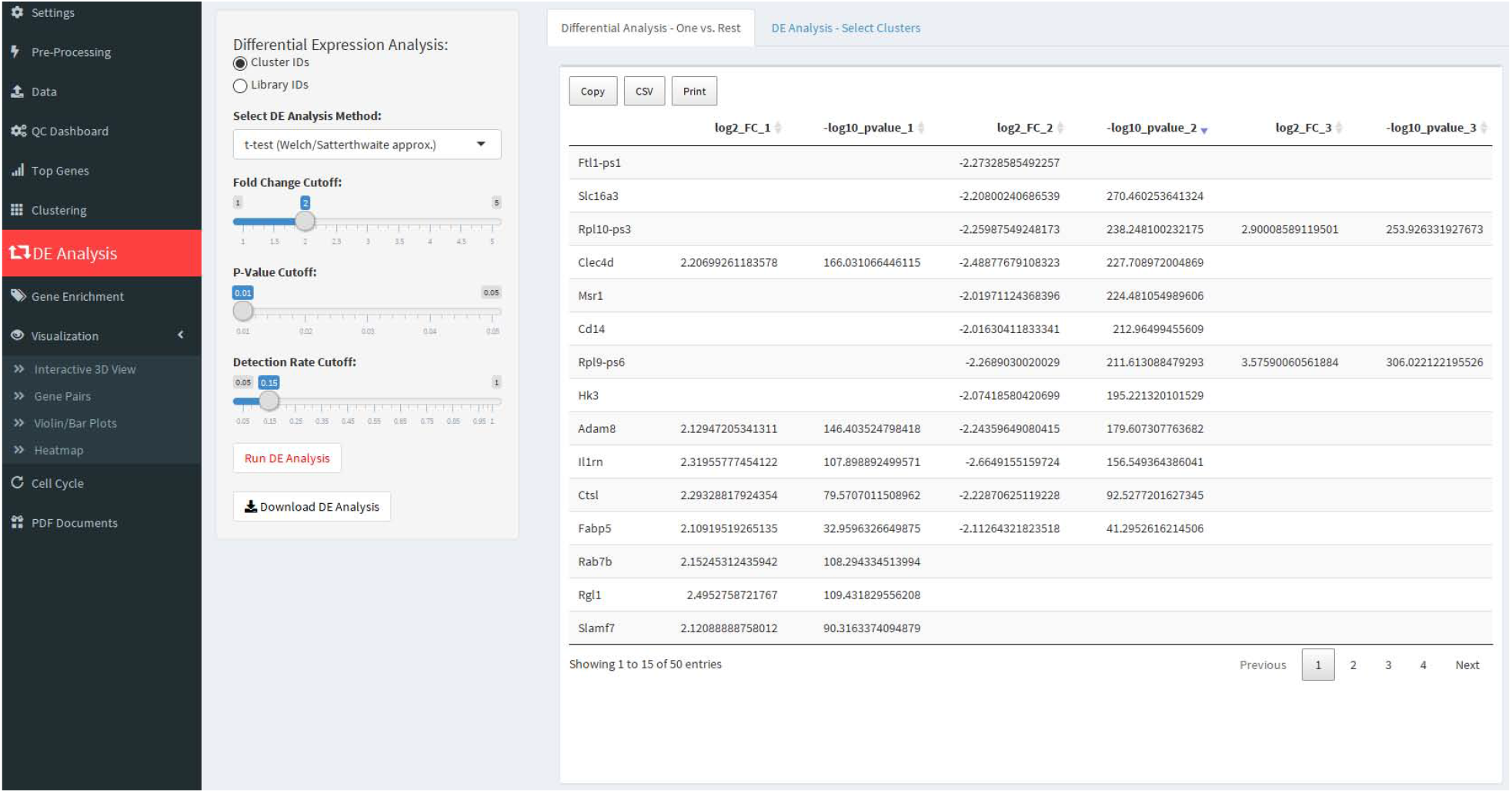
SC1 differential expression analysis.

**Figure 10.**
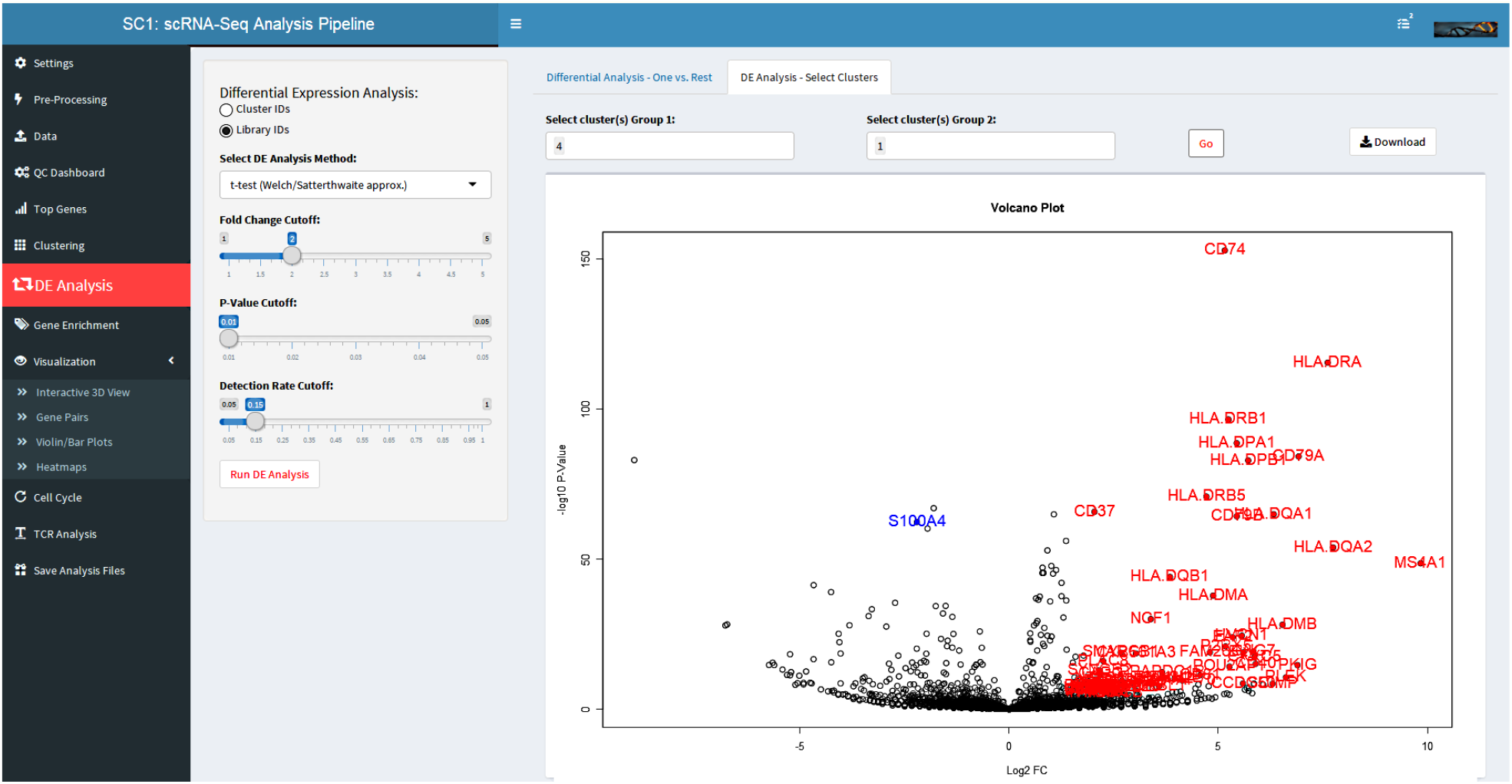
SC1 differential expression analysis between two user-defined cell groups (from the *PBMCs* dataset). Results illustrated as a volcano plot.

#### Enrichment Analysis

DE analysis is followed by cluster-based gene functional enrichment analysis performed using the ‘gProfileR’ and ‘gprofiler2’ R packages Reimand et al. (2007) with results visualized as word clouds (Fig. 11) and provided as downloadable term significance values to help with cluster annotation. Labels assigned to the clusters at this step update throughout SC1 tool output and visualization plots.

**Figure 11.**
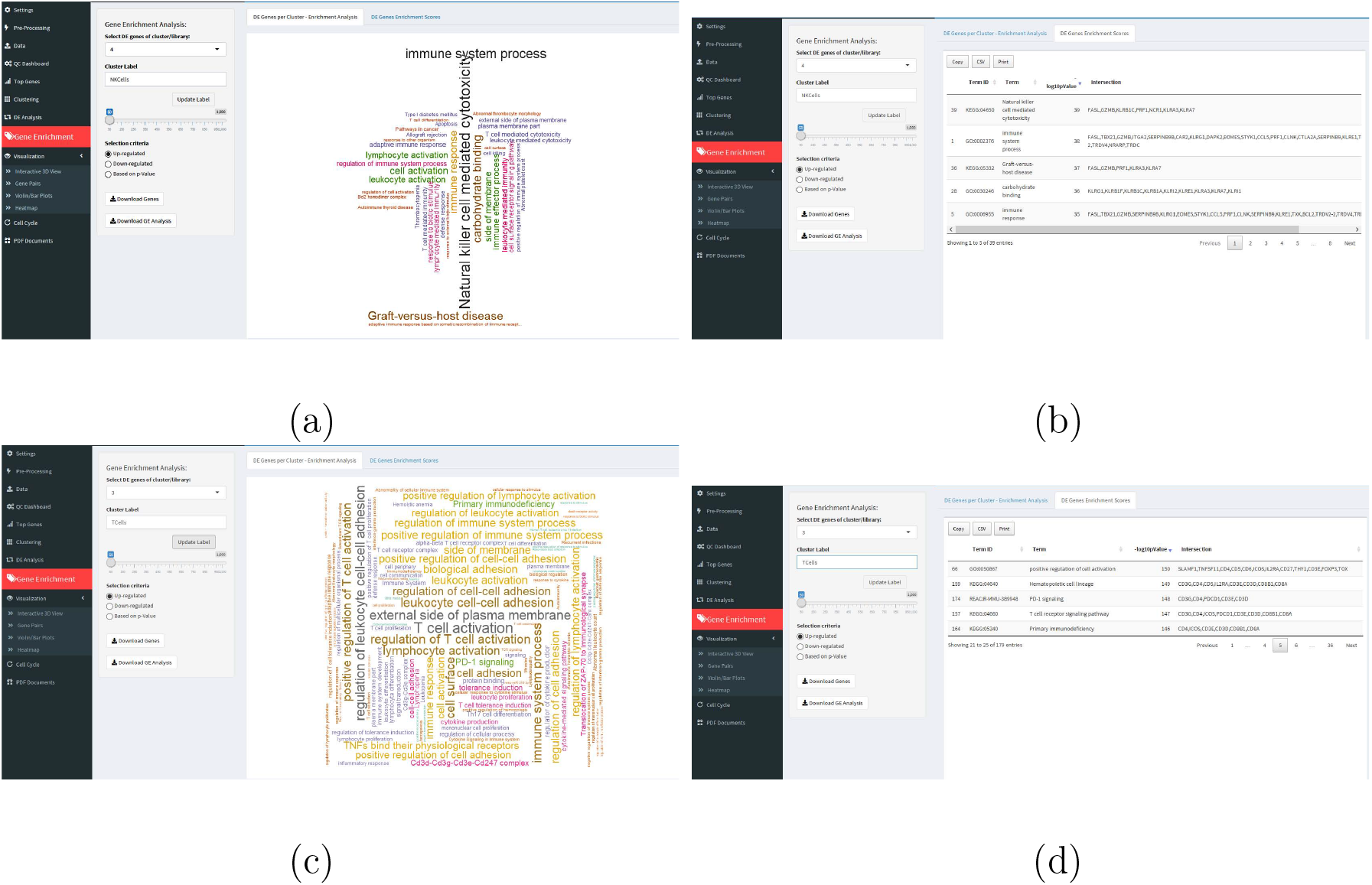
SC1 cluster-based gene enrichment analysis. a) and c) World Cloud visualization of the most significant biological terms identified from the top up-regulated DE genes in a selected cluster. b) and d) Most significantly enriched DE gene modules per selected cluster.

#### Interactive Data Visualization

In previous sections several visualization tools and plots supported in SC1 were described, including the interactive 2D data views in section 2.2, vol-cano plots for differential expression visualization in section 2.2 and word clouds showing gene enrichment in section 2.2. Many other SC1 analysis steps generate visualizations of the results, including for instance the violin plots showing the probability density of gene expression values for each selected cluster/library and the bar-plots showing percentage of cells expressing selected genes by cluster or by library (Fig. 12), also dendrogram plots are used in SC1 to show clustering hierarchies or cell cycle reconstructed order. More visualizations are discussed in the following sections and can be explored through loading any of the provided example datasets into SC1 tool at https://sc1.engr.uconn.edu/.

**Figure 12.**
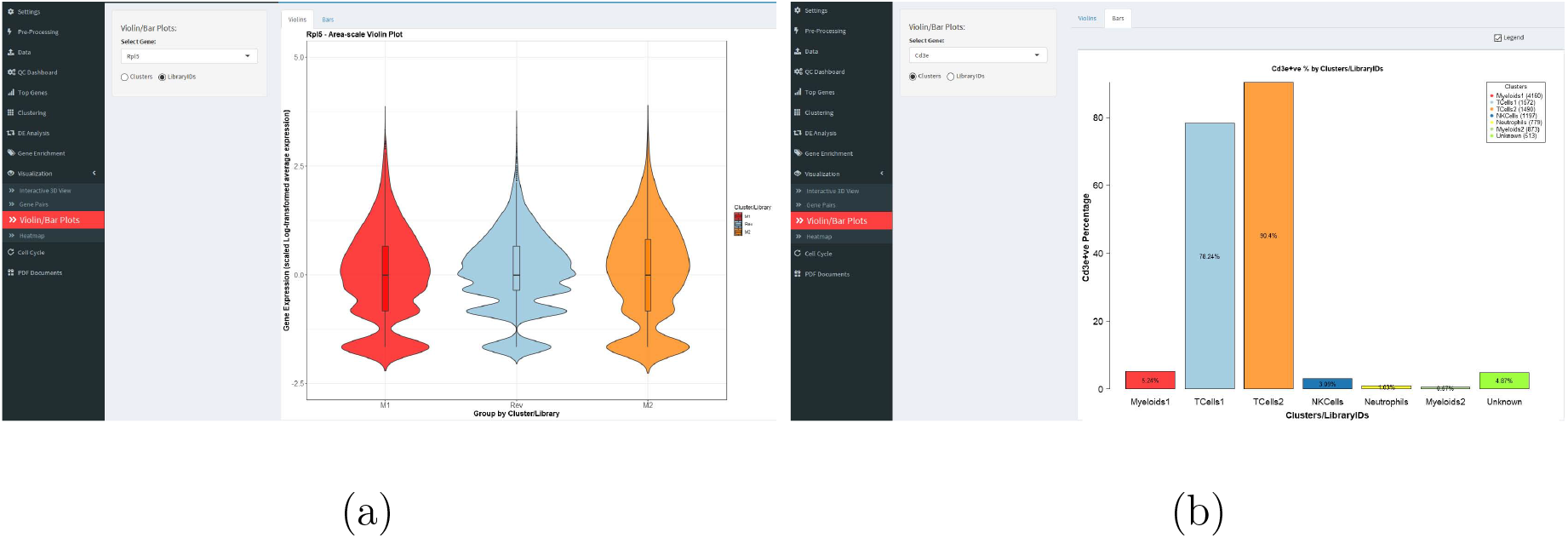
SC1: Violin a) and Bar b) plots by cluster or library/sample.

Additional visualizations include

- Clustering and gene co-expression visualization. SC1 includes multiple interactive visualization options; the interactive 3D t-SNE or UMAP visualization tabs include the ability to select genes individually, in pairs, or in groups as predefined gene sets. Cells are identified where all (AND) or any (OR) of the selected genes are detected. Identified cell populations can be selected or excluded to form a subset that can be downloaded and used to form a new sub-population for further analysis in SC1 (Fig. 13). Identifying various cell populations in SC1 and downloading relevant cells’ expression profiles can be achieved in various ways in SC1: by selecting pre-defined libraries or conditions or selecting cell populations based on gene selection, also selecting specific cell types from clustering analysis results. Gene pair co-expression can be visualized using interactive 3D plots as well as scatter plots as shown in Fig. 14.

**Figure 13.**
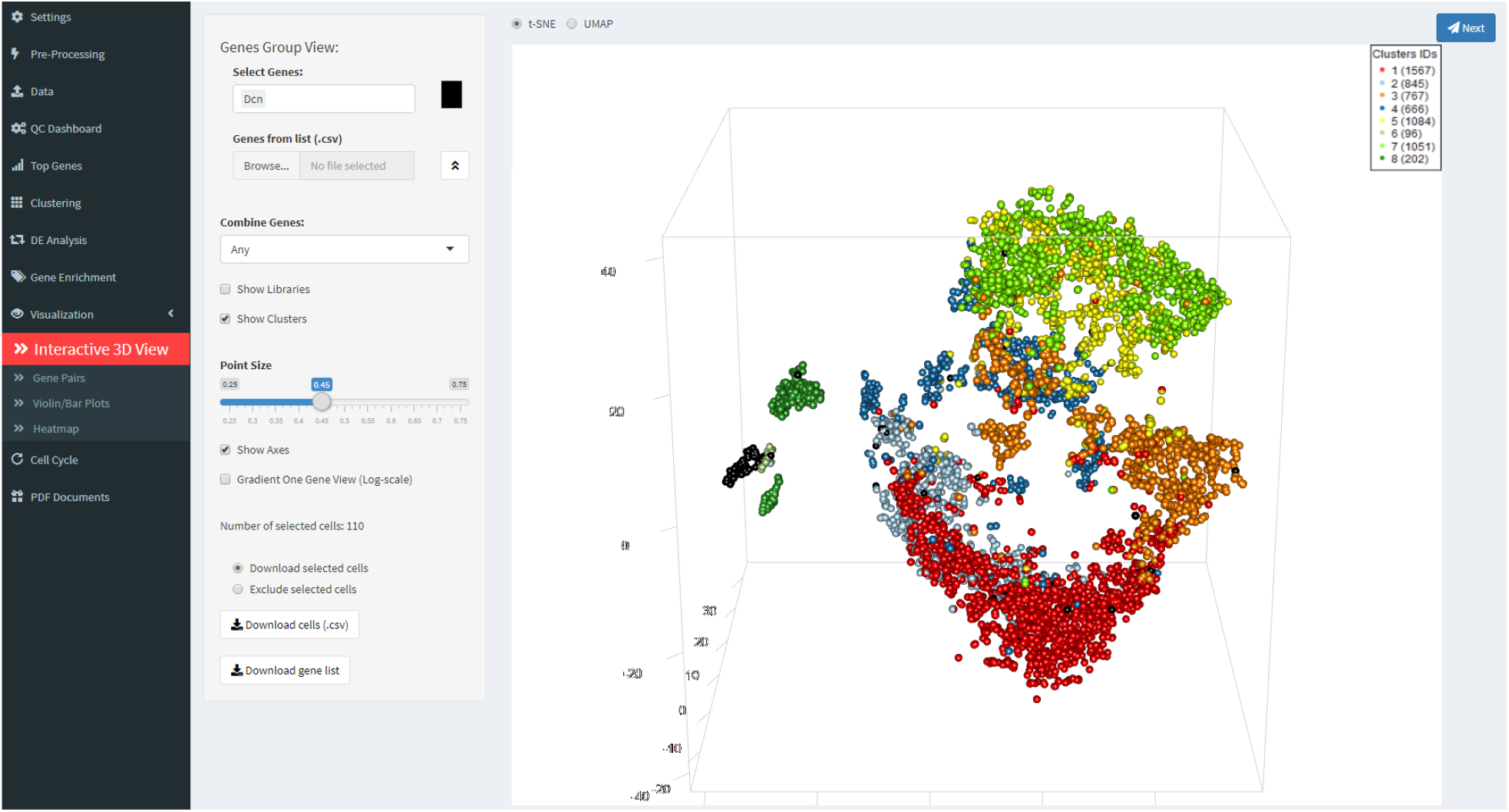
SC1 3D visualization of clustering results and selected genes from the *HPV* data from Lukowski et al. (2018).

**Figure 14.**
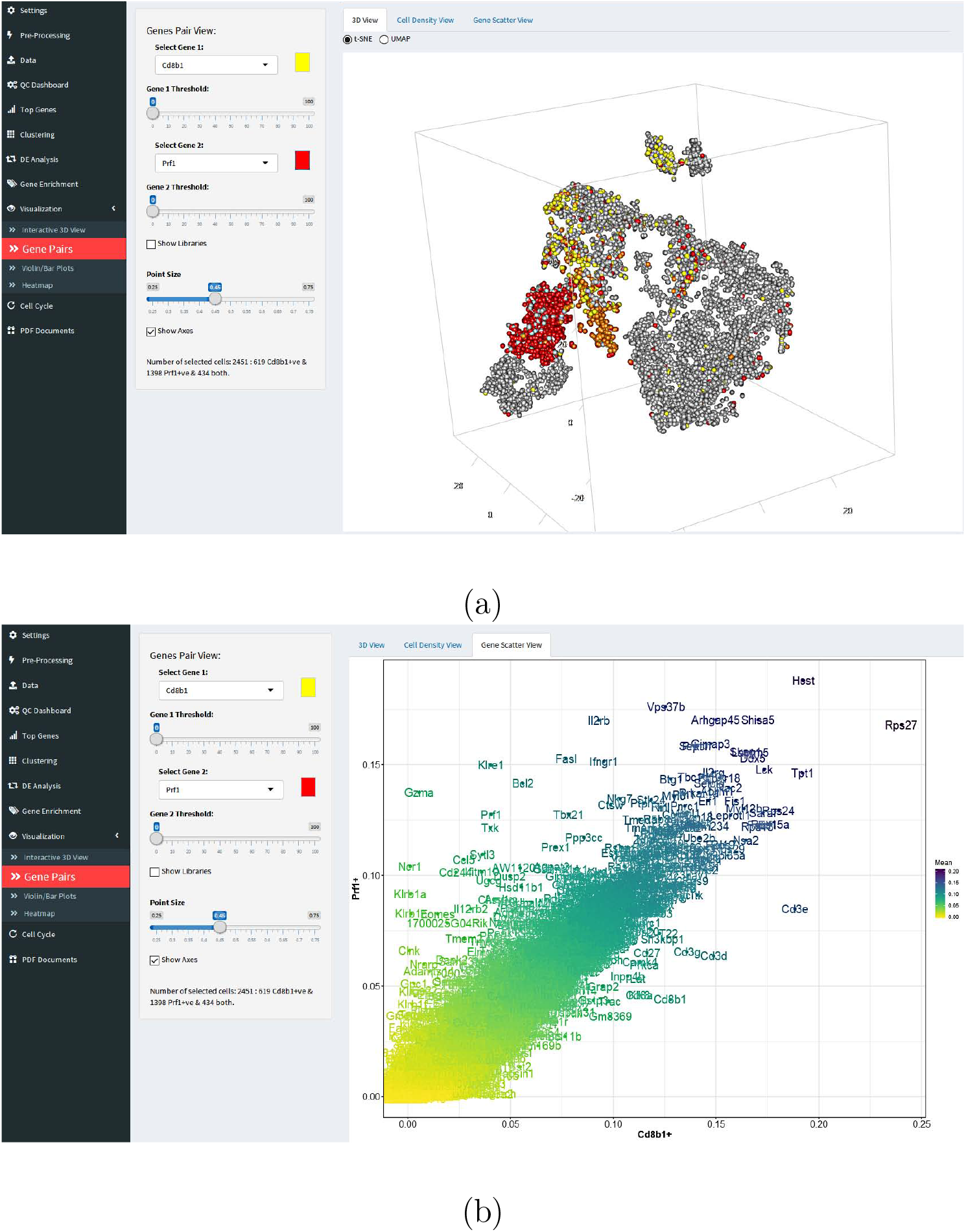
SC1 gene co-expression visualization show for selected Gene-1 and Gene-2 a) 3D tSNE projection highlighting cells expressing each of the selected genes in yellow and red as well as cells expressing both selected genes in orange (auto color mixture) and b) scatter plot contrasting the Gene-1 positive and Gene-2 positive cell populations’ average gene expression values on the x and y axis of the plot. Example from the *HPV* dataset Lukowski et al. (2018).

- Detailed and summary heatmaps. *Detailed Heat Map View:* SC1 provides several ways to select genes and cells visualized in configurable heat maps. Automatic identification of informative genes based on average TF-IDF allows the generation of exploratory heat maps to investigate the heterogeneity of the data. A list of highly expressed/abundant genes can also be downloaded from SC1 and used to construct a heat map. SC1 also supports custom gene selection by manually selecting or uploading a list of genes of interest to use for heat map construction. After the DE analysis step is concluded, the list of differentially expressed genes can also be visualized as a heat map. The expression/count values are by default log transformed in SC1 heat maps using the *log*2(*x* + 1) transformation. *Summary Heat Map View:* The summary heat map view in SC1 provides a “pseudo-bulk” view of the data, showing average expression profiles for selected genes by cluster or library (Fig. 15). The gene expression levels in summary heat maps are row-normalized, i.e., gene means expressions in libraries and clusters are normalized by dividing by the max mean expression of each gene over all libraries and clusters. This assigns a maximum value of 1 (red) to the groups for which the mean expression of the gene is the highest.

**Figure 15.**
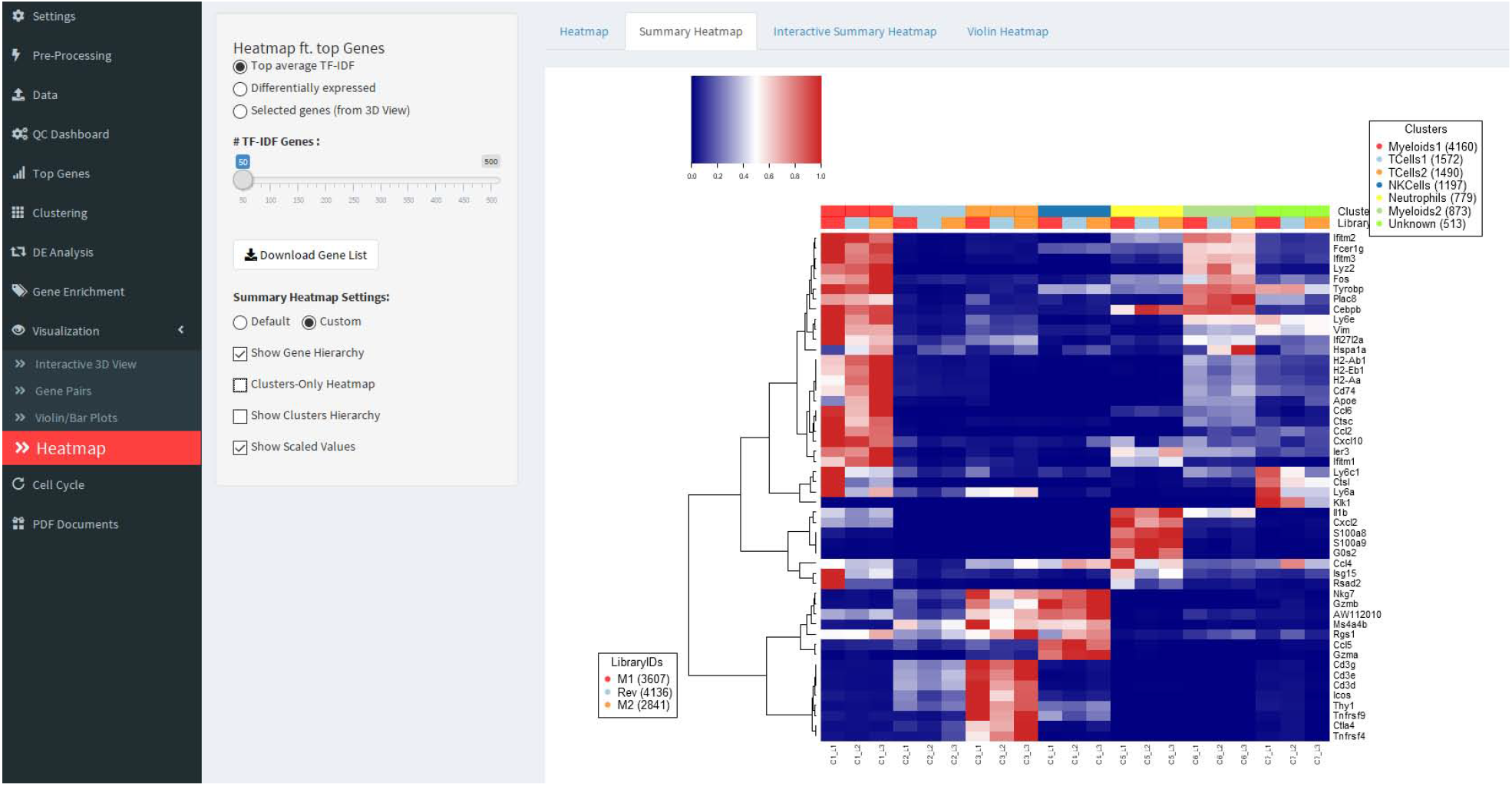
Summary heat map showing cluster/library breakdown mean expression profiles of selected genes.

Other summary heatmap views implements a cluster-based average expression view of selected gene lists (options include DE gene lists, TF-IDF gene lists or user-selected gene lists) as shown in Fig. 16.

**Figure 16.**
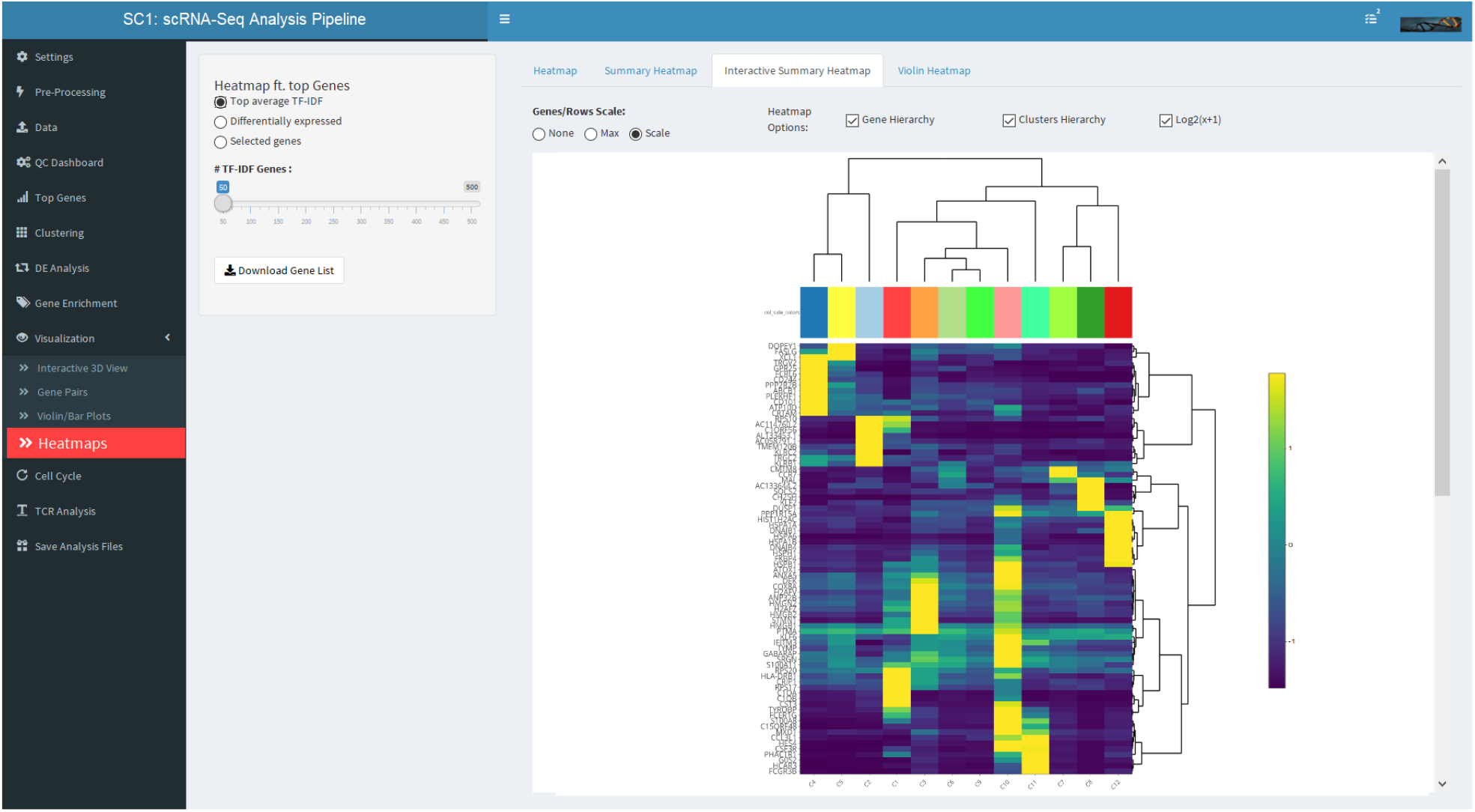
Summary heat map showing cluster based mean expression profiles of T cells of the *COVID19* set for selected DE genes.

#### Cell Cycle Analysis

In SC1, an orthogonal analysis of cell cycle effects can be performed at any stage of the analysis by clustering and ordering cells according to the expression levels of cell cycle genes as detailed in Moussa (2018) and Moussa & Măndoiu (2020a). Here we provide a brief summary since the cell cycle effect, i.e. gene expression profile variation in single cells due to difference in their cell cycle phases, can present a prominent source of variance between cells and can often interfere with cell type or cell state identification and functional analysis of scRNA-Seq data.

SC1 supports three different cell cycle gene lists: genes annotated with the “cell cycle” GO term (GO:0007049) in the Gene Ontology database Consortium (2004), genes from the Cyclebase 3.0 database Santos et al. (2014), and finally the list of periodic genes identified in Dominguez et al. (2016). The list of cell cycle genes can be further filtered based on the gene expression values in the current dataset analyzed in SC1 so as to keep a correlated set of expressed genes with higher than *α* correlation filter threshold, this step can filter out genes that - although annotated as cell cycle related - do not highly correlate with other expressed cell cycle genes in the current dataset, and can hence be considered outliers. SC1 implementation of the method from Moussa & Măndoiu (2020a) auto-determines the number of PCs to use in this analysis by assessing the drop in variance explained for each pair of consecutive principal components. SC1 users can also manually specify the number of PCs if desired. The principal component analysis is followed by a 3-D t-SNE transformation. T-SNE based dimensionality reduction using the main principal components captures local similarity of the cells without sacrificing the global variation already captured by the principal component analysis. Cells are subsequently clustered into a hierarchical structure based on their expressed cell cycle genes profile similarity. Cosine similarity is used as the default similarity metric.

The cell cycle is typically divided into 6 distinct phases (G1, G1/S, S, G2, G2/M, and M, Cooper et al. (2000)), a default of up to 7 clusters can be identified from the hierarchical clustering corresponding to the 6 cell cycle phases plus at least one potential cluster of non-cycling cells (Fig. 17). The maximum number of clusters can be manually modified by the user in SC1 and an ‘auto’ option for determining the optimal number of clusters based on the Gap Statistics Analysis algorithm from Tibshirani et al. (2001) is also implemented in SC1.

**Figure 17.**
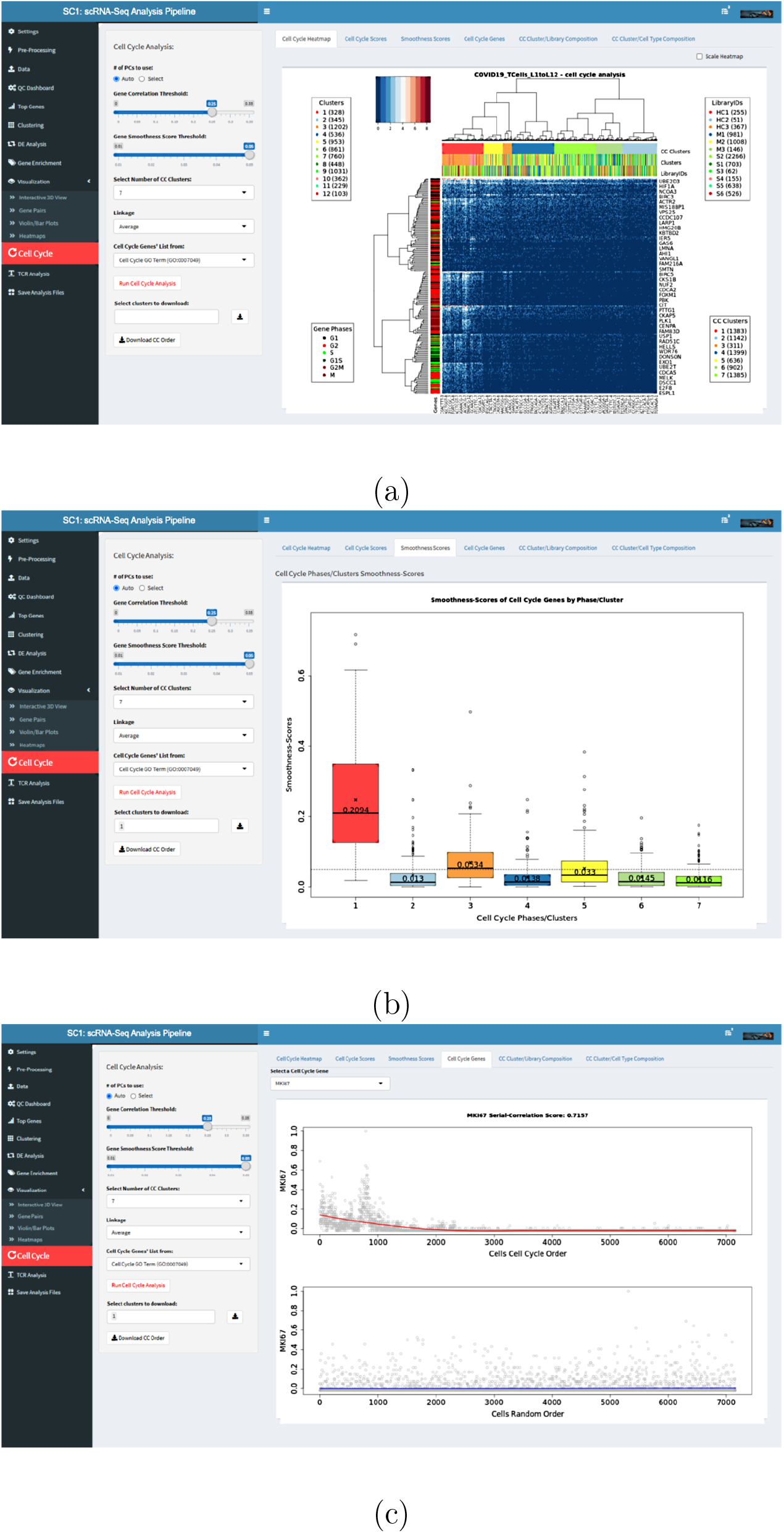
*COVID19* dataset cell cycle analysis in SC1. a) Heatmap showing CC genes and clusters. b) Gene Smoothness Scores per CC cluster. c) Cell cycle gene expression ordered by identified CC order.

An order of cells reflecting their position in the progression along the cell cycle phases is reconstructed by reordering the leaves of the hierarchical clustering dendogram using the *Optimal Leaf Ordering (OLO)* algorithm Bar-Joseph et al. (2001) as implemented in Hahsler et al. (2008). For more details on this reconstruction the user is referred to Moussa & Măndoiu (2020a)

#### Cell Cycle Cluster Scores

In SC1 implementation of the cell cycle analysis approach, two types of scores are calculated for each identified cell cycle cluster, Cluster Mean-Scores and Gene-Smoothness Score (GSS). For details about the two metric, we refer the reader to Moussa & Măndoiu (2020a).

In summary, the Cluster Mean-Score determines the cell cycle phase: 6 gene groups (G1, G1/S, S, G2, G2/M, and M genes, respectively) which include cell cycle genes that are known to reach their peak expression in the corresponding cell cycle phases Liu et al. (2017) are evaluated per cluster. For each of these genes and each cell, a ‘z-score’ is computed by subtracting the gene’s mean expression level from the expression level of the gene in the cell and then dividing by the gene’s standard deviation. For each group of genes and each cluster identified during the hierarchical clustering step we compute a mean-score by averaging over cells in the cluster and genes in the group. The maximum mean-score of a cluster is used to determine its cell cycle phase.

Gene-Smoothness Score is an independent metric that can be used to distinguish dividing from non-dividing cells. Normalized gene scores computed as the described Cluster Mean-Scores are relative between cell cycle phases and cannot always clearly distinguish, between dividing vs. non-dividing cells. The GSS can therefore be used for this purpose, as well as help assess the cells order according to their cell cycle phase progression. The GSS is based on serial correlation, the correlation between a given variable and a lagged version of itself and can be computed for any ordered group of cells to directly assess the suggested order. Strengths of this metric include the fact that the cells do not need to have known cell-cycle labels and that no specific model assumptions are required for the marker gene expression (whether binary, bi-modal, sinusoidal, etc.).

The GSS of an ordered cluster/group *c* of cells is defined in Moussa & Măndoiu (2020a) as

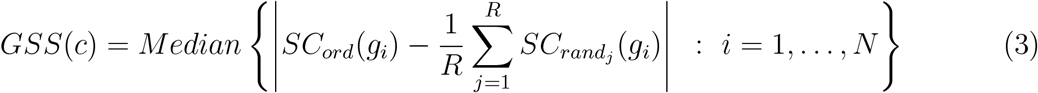

where *N* is the number of annotated cell cycle genes, *SC*_*ord*_(*g*_*i*_) denotes the first-order serial correlation of gene *i* with respect to the given cell order, and *SC*_*rand*_*j* (*g*_*i*_), *j* = 1, …, *R*, denote the first-order serial correlation of gene *i* with respect to *R* randomized cell orders.

The first-order serial (or auto-) correlation is the correlation value between a given gene expression vector and a version of itself shifted by one position. Serial correlation is a value between −1 and 1. First-order serial correlation near 0 implies that there is no overall correlation between adjacent data points. On the other hand, a first-order serial correlation near 1 suggests a smoothly varying series, while a first-order serial correlations near -−1 indicates a series that alternates between high and low values. Because individual cell cycle genes can be expressed in different patterns throughout the cell cycle phase transitions, and even abruptly switch direction when the assessed cluster includes mostly cells in one of the transient cell cycle phases (G1/S or G2/M), GSS is defined as the median (over all cell cycle genes) of the absolute differences between the serial correlation of a gene’s expression values ordered according to the given cell ordering and the average serial correlation computed over *R* randomized orders.

A cluster of cells is considered to be dividing when its GSS is greater than a certain error margin *ε* (default 0.05), translating to at least 50% of the genes have an absolute difference in serial correlations between randomized order and identified cell cycle order of at least 0.05. SC1 implementation allows the user to select any cell cycle gene of interest and examine its normalized expression levels along the inferred order as seen in Fig. 17c), where gray dots represent normalized gene expression values for individual cells, while the red and blue curves represent the fitted local polynomial regression of these values for the SC1CC and a random cell order, respectively. The fitted expression lines under random ordering of the cells convey no recognizable pattern and stay nearly flat close to an altitude of 0 and in contrast, the identified cell order in SC1 results in fitted curves that appear to peak at different positions, consistent with these gene’s involvement in different cell cycle phases.

#### TCR-Seq Analysis

SC1 integrates single cell TCR-Seq data and links clonotype data to clusters identified from the scRNA-Seq gene expression data analysis (this analysis step is compatible with TCR-Seq data from 10x Genomics). TCR-Seq clonotypes stacked bar plot in Fig. 18 shows the TCR-Seq data from Brennick et al. (2021) illustrating clonotypes by cluster, each segment of the stacked bar represents a unique clonotype with height representing the number of clones, the total height represents the total number of clones per cluster.

**Figure 18.**
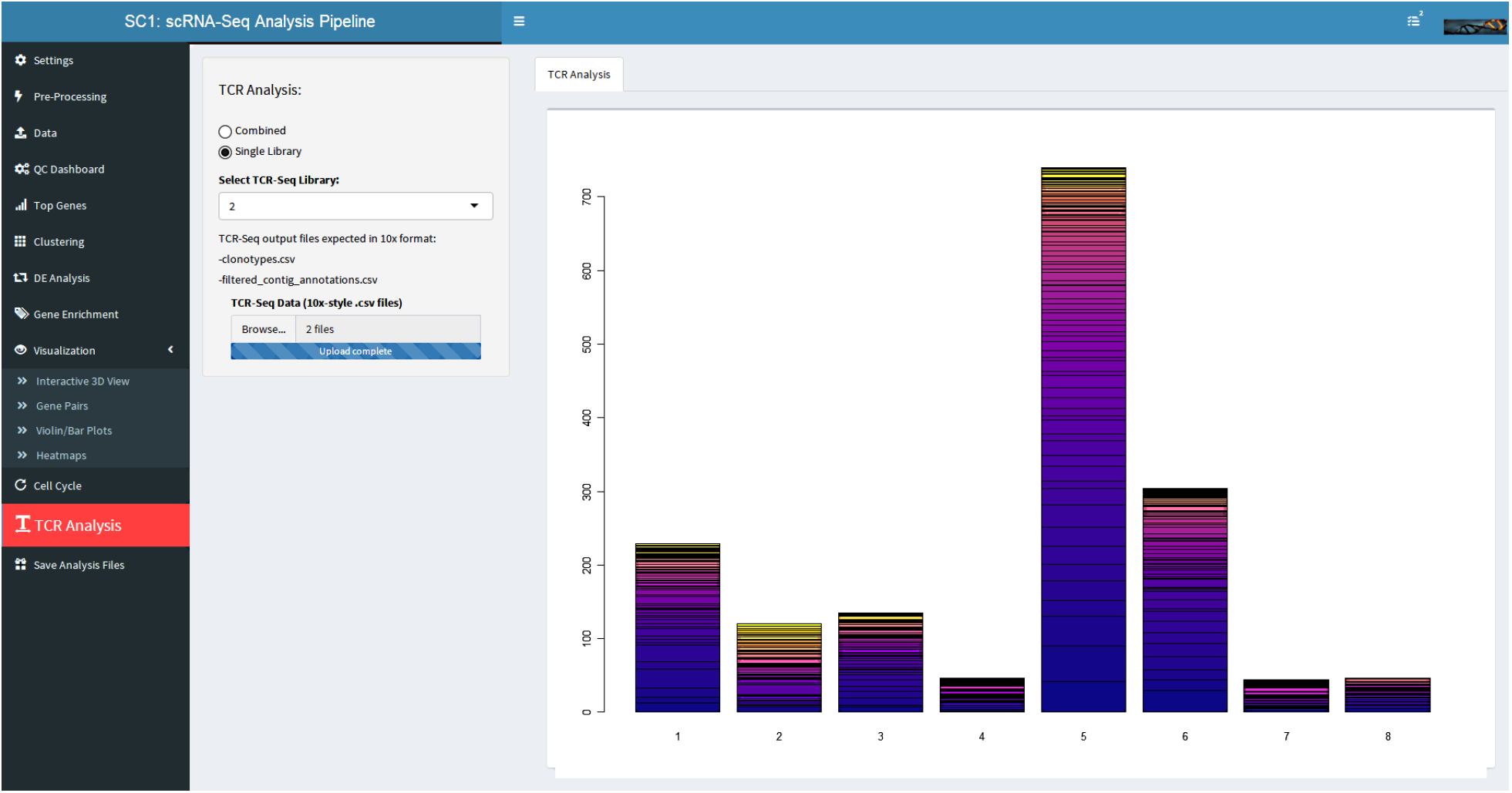
TCR-Seq clonotypes stacked bar plot showing clonotypes by cluster, each segment of the stacked bar represents a unique clonotype with height representing the number of clones, the total height represents the total number of clones per cluster.

## 3 SC1 Applications in Immunology

### *SNS* dataset analaysis

**Nevin et al. (2020)** In a recent study from Nevin et al. (2020) the characteristics of the myeloid compartment of SNS-ablated tumor-bearing mice were investigated by single cell RNA sequencing. In this section we show how SC1 pipeline can be used to fully characterize the scRNA-Seq dataset from this study (GEO accession number GSE154973). Sorted CD3-CD19-cells from the spleen of SNS-ablated or vehicle control treated CT26 tumor-bearing mice and captured using the 10x sequencing are first clustered into a myeloid and natural killer cells compartments. One of the goals of this part of the analysis is to examine the myeloid compartment in detail, so the natural killer cells population identified at the first tier of the dataset analysis in SC1 is excluded and the rest of the cells population are downloaded from SC1 and re-processed as a new sub-dataset (Fig. 19).

**Figure 19.**
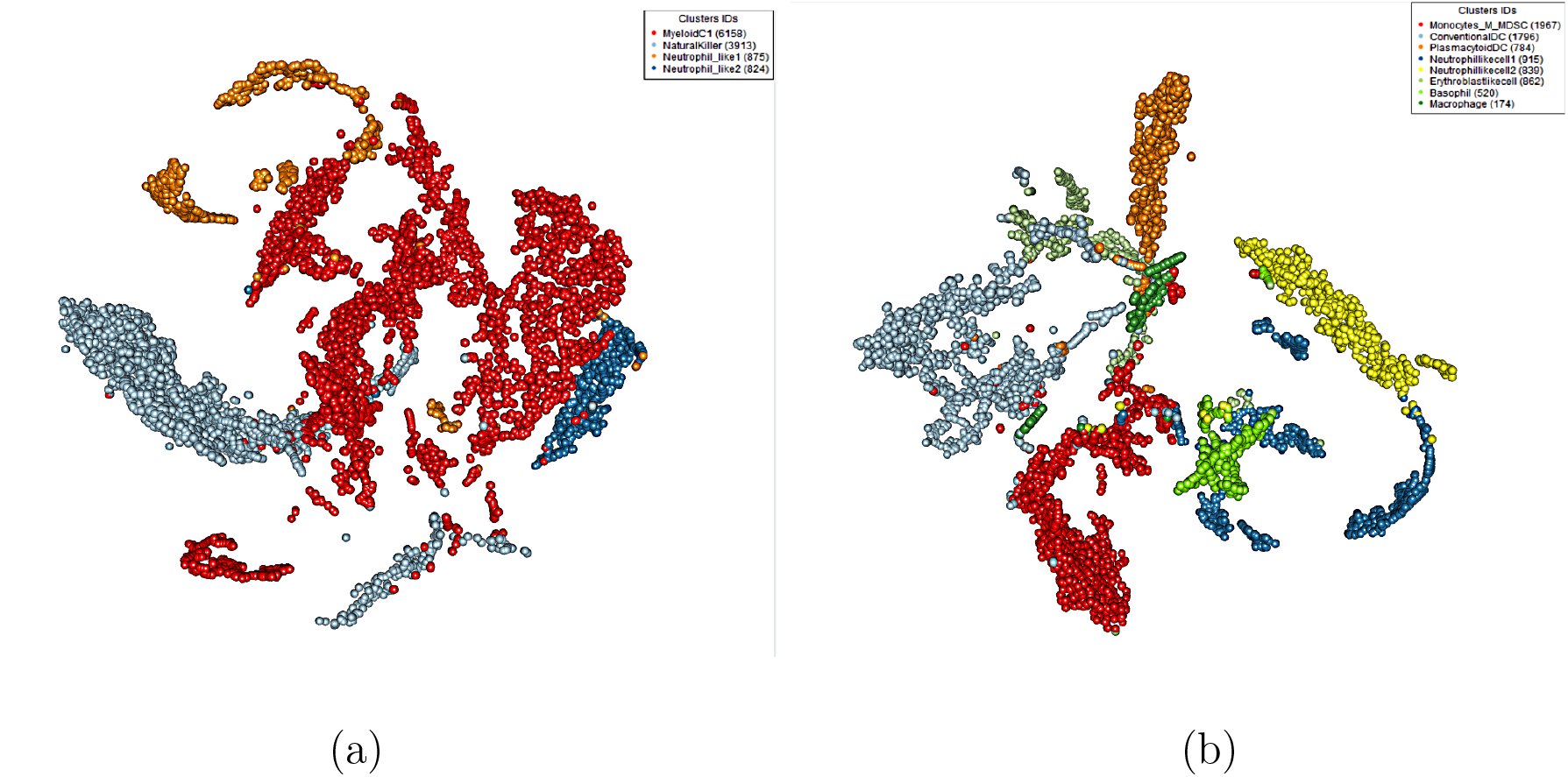
3D tSNE projection of the *SNS* dataset initial clustering into 4 main populations (a) followed by the exclusion of the Natural Killer cells population and re-clustering (b).

The myeloid compartment clustered into eight distinct populations with two distinct clusters of polymorphonuclear neutrophil (PMN)–like cells are identified (>99% of cells in these clusters expressing both S100a8 and S100a9). These two clusters in turn are further reanalyzed, and characterized into five distinct populations (Fig. 20). At this second tier of the analysis, clear expansion of PMN cluster 4 was observed from 10.71 to 21.01% of PMNs upon SNS ablation in tumor-bearing mice. This cluster was defined by relatively high expression of S100a8 and S100a9, high expression of Cybb (Nox2), differential up-regulation of transcription factors Stat3 and Cebpb, and differential up-regulation of the suppressive cytokine Tgfb1 (Fig. 20) all of which are associated with the PMN-MDSC phenotype Nevin et al. (2020).

**Figure 20.**
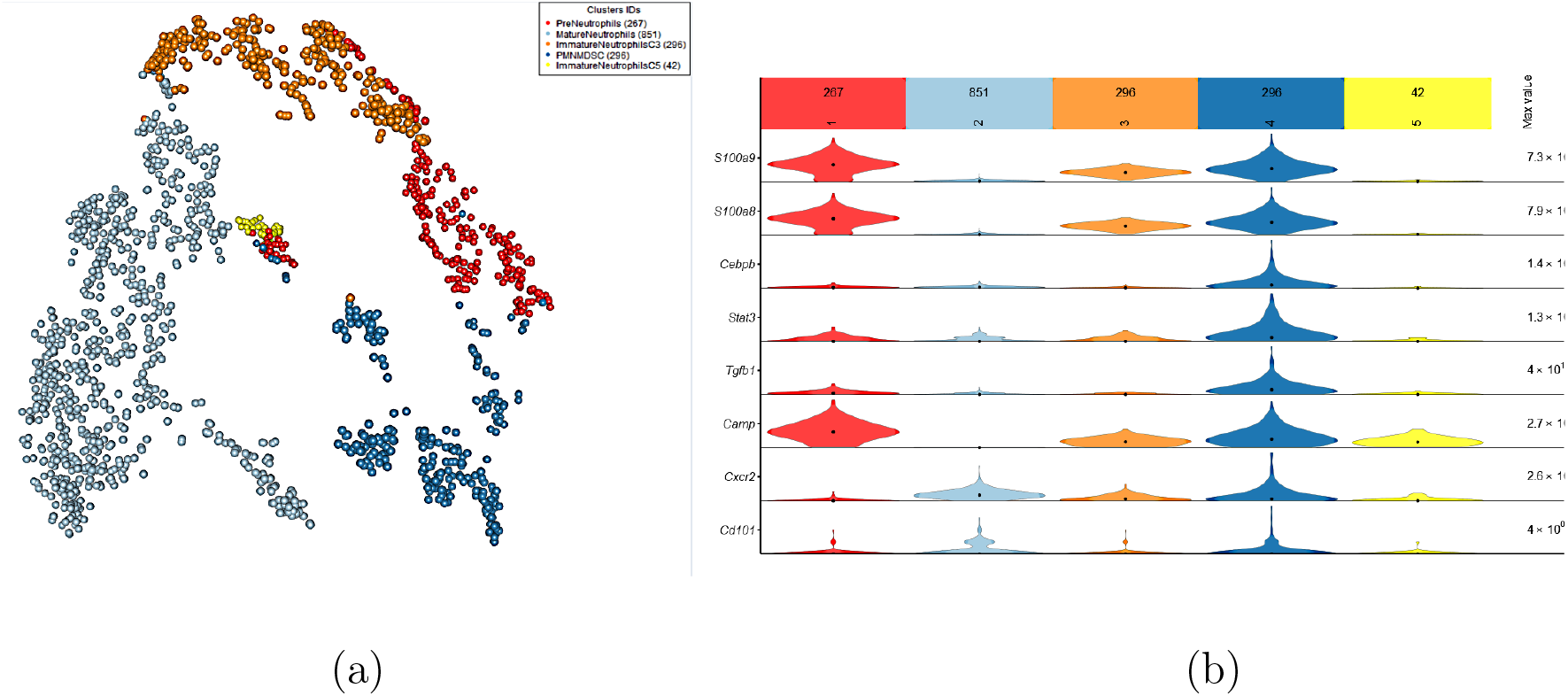
Sympathetic nervous tone limits the development of myeloid-derived suppressor cell. 3D tSNE projection of the neutrophil-like population of the *SNS* dataset, annotated according to the DE analysis. Select DE genes expression shown in violin heatmap (AllenInstitute (2018)).

## 4 Conclusion

SC1 provides a powerful tool for interactive web-based analysis of scRNA-Seq data. The SC1 workflow is implemented in the R programming language, with an interactive web-based front-end built using the Shiny framework Chang et al. (2017). SC1 employs a novel method for gene selection based on *Term-Frequency Inverse-Document-Frequency (TF-IDF)* scores Moussa & Măndoiu (2018), and provides a broad range of methods for cell clustering, differential expression analysis, gene enrichment, visualization, and cell cycle analysis. Future work includes integrating additional clustering methods, as well as other differential expression analysis methods and integrating methods for cell differentiation analysis. As the amount of scRNA-Seq data continues to grow at an accelerated pace, we hope that SC1 will help researchers to fully leverage the power of this technology to gain novel biological insights.

